# BIRD: Identifying Cell Doublets via Biallelic Expression from Single cells

**DOI:** 10.1101/709451

**Authors:** Kerem Wainer-Katsir, Michal Linial

## Abstract

**Motivation:** Current technologies for single-cell transcriptomics allow thousands of cells to be analyzed in a single experiment. The increased scale of these methods led to a higher risk of cell doublets’ contamination. Available tools and algorithms for identifying doublets and estimating their occurrence in single-cell expression data focus on cell doublets from different species, cell types or individuals.

**Results:** In this study, we analyze transcriptomic data from single cells having an identical genetic background. We claim that the ratio of monoallelic to biallelic expression provides a discriminating power towards doublets’ identification. We present a pipeline called BIRD (BIallelic Ratio for Doublets) that relies on heterologous genetic variations extracted from single-cell RNA-seq (scRNA-seq). For each dataset, doublets were artificially created from the actual data and used to train a predictive model. BIRD was applied on Smart-Seq data from 163 primary fibroblasts. The model achieved 100% accuracy in annotating the randomly simulated doublets. Bonafide doublets from female-origin fibroblasts were verified by the unexpected biallelic expression from X-chromosome. Data from 10X Genomics microfluidics of peripheral blood cells analyzed by BIRD achieved in average 83% (± 3.7%) accuracy with an area under the curve of 0.88 (± 0.04) for a collection of ∼13,300 single cells.

**Conclusions:** BIRD addresses instances of doublets which were formed from cell mixtures of identical genetic background and cell identity. Maximal performance is achieved with high coverage data. Success in identifying doublets is data specific which varies according to the experimental methodology, genomic diversity between haplotypes, sequence coverage, and depth.

## 1 INTRODUCTION

Single-cell RNA sequencing (scRNA-seq) technology has evolved very rapidly in recent years (Kolodziejczyk, et al., 2015; Lan, et al., 2017; Zheng, et al., 2017; Zilionis, et al., 2017). scRNA-seq enables higher resolution of the expression profile of cells within cells tissue and enables accurate assessment of the single cells’ identity and variability. This new technology has been applied to a wide range of biological studies and across many organisms, including rodents and humans. Complex tissues were dissociated and sequenced by scRNA-seq resulting in cataloging cells by their types determining tissue composition (Usoskin, et al., 2015; Villani, et al., 2017; Zeisel, et al., 2015) and identifying overlooked cell types (Buettner, et al., 2015).

All scRNA-seq studies rely on profiling cell transcriptomes. The main hurdle in obtaining reliable and high-quality data from scRNA-seq stems from the limited amounts of RNA per cell and the stochastic nature of transcription (Ilicic, et al., 2016). Specifically, the majority of current scRNA-seq methods suffer from low capture efficiency and high dropouts (Haque, et al., 2017). Additionally, all single-cell expression data are signified by a strong signal of monoallelic expression which is not detected from the sequencing pools of cells. The dominant monoallelic expression of single cells (Borel, et al., 2015; Jiang, et al., 2017) is attributed to allelic dropout of transcripts due to the insufficient coverage, and to the cellular phenomenon of “transcriptional burst” (Reinius and Sandberg, 2015). The latter means that each of the alleles has its kinetics, thus at any specific time expression mostly stems from a single allele (Larsson, et al., 2019).

Innovative technologies for scRNA-seq were developed to increase high throughput while minimizing biological intrinsic and technical errors (discussed in (Bacher and Kendziorski, 2016; Chen, et al., 2019; Hashimshony, et al., 2016)). Some methods make use of fluorescence-activated cell sorting (FACS) (Kolodziejczyk, et al., 2015; Wagner, et al., 2018) and microfluidic-based platforms, such as the C1 Single-Cell Auto Prep System (Fluidigm) (Xin, et al., 2016). These methods are usually followed by a full-length transcript sequencing as in Smart-seq2 (Picelli, et al., 2014). Others use droplet microfluidic procedures that combine a tagging step before cell lysis (reviewed in (Chen, et al., 2019; Klein, et al., 2015). Advances in the droplet technique allow capturing beads with a single cell per droplet (dscRNA-seq) thus increasing the scale for single-cell transcriptomic by two orders of magnitude (Fan, et al., 2015; Sheng, et al., 2017). These protocols primarily use only poly-A sequencing and are thus biased toward the 3’ side of the transcript. Most current-day protocols include additional steps of barcoding the transcripts by UMIs (unique molecular tag identifiers) (Klein, et al., 2015), and further improvement of the capturing efficiency (Sheng and Zong, 2019).

One of the pitfalls in the field concern a faulty identification of a doublet of cells as a single cell. Doublets rate depends on the concentration of the input cells which is estimated from the dilution Poisson statistics (Macaulay, et al., 2017). An increase in doublets rate is also associated with the unique features of the subjected tissue and cells’ isolation protocols. New methods for increasing cell capturing that reduces costs include multiplexing protocols (Zheng, et al., 2017). By increasing the number of cells as input, the multiplexed droplet RNA-Seq (dscRNA-seq) benefits from reducing technical noise (Zhang, et al., 2019). However, as a byproduct, it leads to an unavoidable increase in the number of cell doublets. One of the methods to identify the rate of doublets in the data includes mixing cells from different origin (e.g., rodents and human (Zheng, et al., 2017)). Alternatively, the dscRNA-seq setting was carried over single cells from several individuals with a different genomic background that were intentionally mixed for estimating the fraction of doublets in the sample (Kang, et al., 2018). Benefiting from the SNP profile of each individual, the Demuxlet algorithm was applied to estimate mixed individual doublets (Kang, et al., 2018). A recently published Scrublet algorithm analyzes single cells for identifying problematic multiplets according to the nearest neighbor graph-based classifier (Wolock, et al., 2019). DoubletFinder (McGinnis, et al., 2019) makes use of the unique cell-state expression profiles for identifying doublets from transcriptionally distinct cells. While these set of methods can differentiate cell mixtures from distinct individuals and cell types, they do not attempt to differentiate cells that originate from the same source or cell type.

In this study, we analyze data from scRNA-seq and dscRNA-seq for identifying doublets without any prior knowledge of cell-type composition. Instead, we take advantage of monitoring allele-specific expression biases. The method called BIRD (BIallelic Ratio for Doublets) relies on analyzing heterologous SNPs present in scRNA-seq data. We report on the accuracy of identifying doublets which is strongly dependent on sequencing methodologies, coverage, depth and the degree of allelic diversity in the genomic data.

## 2 MATERIALS AND METHODS

### 2.1 Dataset of single cells

#### Dataset 1. Primary human fibroblasts

A dataset of scRNA-seq of female fibroblast UCF1014 was downloaded from the European Genome-phenome Archive (https://www.ebi.ac.uk/ega/home) using accession number EGAD00001001083. The data consist of two sets of scRNA-seq: 104 cells (22 PCR cycles) and 59 cells (12 PCR cycles). The data was collected in a C1 Auto Prep System (Fluidigm) device and sequenced using full transcript Smart-seq2 (Picelli, et al., 2014). DNA-seq of UCF1014 was also downloaded from EGAD00001001084. The sequence data was produced and described by (Borel, et al., 2015).

#### Dataset 2: Peripheral human blood mononuclear cells

The data was created and described in (Kang, et al., 2018). Peripheral blood mononuclear cells (PBMCs) scRNA-seq from 8 different individuals were downloaded from the Gene Expression Omnibus (GEO) database, accession number GSE96583. This dataset contains 3 different runs. Two of the runs include a mixture of scRNA-seq from 4 different individuals (run_a and run_b sets). The third run is a mixture of all 8 individuals scRNA-seq data (run_c). Cells were sequenced using 10X Genomics (Chromium instrument) methodology. Additional VCF files of exome sequencing of these individuals were extracted through Github link (https://github.com/yelabucsf/demuxlet_paper_code/tree/master/fig2). It shares also an additional file determining the individuals’ origin per each scRNA-seq as processed by the Demuxlet tool (Kang, et al., 2018). Only cells that were assigned by Demuxlet to belong to the same individual and therefore could not be explicitly annotated as singlets or doublets were used for further analysis by our methodology.

### 2.2 Biallelic score for single cells

To correctly estimate the Allelic Specific Expression (ASE) and specifically, the degree of biallelic expression of each cell, the DNA-seq of cell line UCF1014 was used to create a collection of all heterozygous SNPs (hSNPs) using Gene Analysis Toolkit (GATK (Van der Auwera, et al., 2013)). All the hSNPs were kept in a VCF file. The RNA-Seq reads were preprocessed using Trimmnomatic (Bolger, et al., 2014) with its default parameters. Using STAR (Dobin and Gingeras, 2015) each scRNA-seq FastQ file was aligned against the GRCh37 (hg19) UCSC female reference (after excluding Y chromosome). The BAM output of the alignment and the hSNPs VCF were processed using Allelcounter-master (Castel, et al., 2015). The tool creates a table containing the number of reads for each SNP that matches the Reference (Ref) and the Alternative (Alt) alleles. Then we processed the table into two tables, one for the Ref alleles and the other for the Alt alleles. Both tables contain the number of reads assigned to each cell for each hSNP. An observation was considered for cells having ≥6 reads for a specific hSNP in a specific cell. The same procedure was applied to all single cell datasets analyzed (described in 2.1).

For the PMBC dscRNA-seq data (Dataset 2) BAM files of the 3 runs were split to single cell BAM files to maintain an individual-based BAM file per cell. Each single cell was identified according to its unique cell-based barcode. Each BAM file was coupled to its corresponding individual VCF according to the identification by the Demuxlet algorithm. (Kang, et al., 2018) and was preprocessed by Allelcounter-master (Castel, et al., 2015). We then unified all cells that share a specific run, for a specific individual into two tables containing the number of reads for each hSNP that matches the Ref and the Alt alleles. As each individual contains its own set of hSNPs, tables for the Ref, Alt were created for each of the 16 run-individual pairs. In this analysis, for an hSNP to be considered, we required the number of reads to be ≥3 per hSNP of the subjected cell.

For both datasets we calculate for every available hSNP the Allelic Ratio (AR) of that hSNP in a specific cell as:

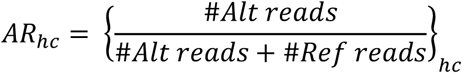

Where h refers to hSNP and c to a specific cell.

The AR ranges from 0 to 1. The value is zero when there was no observed expression from that hSNP. Values of 0.001 and 1 correspond to reads that were fully aligned to the Ref or Alt alleles respectfully. Genuine biallelic hSNP values are bounded by 0.1<=AR<0.9.

An allele independent score for Biallelic Ratio (denoted BAR) was calculated as follows: For a given cell and a given gene, let *i* be an index of the informative (heterozygous) variants, and define by *Ref*_*i*_ and *Alt*_*i*_ the number of reference and alternative reads each informative variant. Define by *Tot*_*i*_=*Ref*_*i*_ +*Alt*_*i*_ the total number of reads for the variant and by *Min*_*i*_=*min* {*Ref*_*i*_, *Alt*_*i*_} the minimal number of reads out of the two alleles of the variant. Let *i*_∗_=*argmax*_*i*_(*Min*_*i*_) be the most informative variant with the maximal biallelic ratio (for the given cell and gene combination). We then define the Biallelic Ratio (BAR) of the cell-gene as:

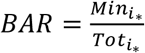

Then, for each cell we take the average BAR of all its expressed genes. Supplemental Fig. S1 shows examples for calculating BAR which ranges between 0 and 0.5. In a formal notation

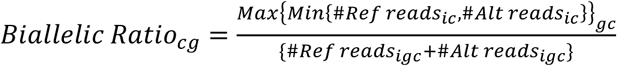

*i*- Heterozygous SNP location (In the numerator stands for the specific SNP that was Max in the denominator), *c* stands for cell and g for a gene.

### 2.3 Doublet simulation and validation

To create a reference dataset of doublets, we created doublets in silico for each of the analyzed datasets separately (see 2.1). For the simulations we randomly sample 10% of the single cells to become cell doublets (creating a composed collection with 5% of the original cells being simulated doublets). We create pairs of cells, and for each pair we sum their corresponding reads from the Ref and Alt tables. Following summation, for the fibroblast data (Dataset 1), we randomly down-sample the reads to the average cell reads number. Due to the low coverage of the PMBCs data (Dataset 2) we skipped this step. In each simulation, we recorded the BAR values for the singlets and the simulated doublets. The procedure of creating simulated doublets was repeated 100 times. For each run, we record the average of the BAR values for all the singlets or all simulated doublets.

The fibroblasts that composed Dataset 1 is of a female origin (Borel, et al., 2015). In these primary cells the Chromosome X inactivation phenomenon was fully maintained (Wainer-Katsir and Linial, 2019). Thus, we used the unique property of X-inactivation to obtain an expression pattern that matches the cell specific activation of one of the X-chromosomes (Garieri, et al., 2018). Specifically, we calculate AR per hSNP per cell. Then, we calculated AR* for assessing the biallelic ratio for chromosome X.

AR* balances between the hSNP expression from either the Ref or Alt alleles. We considered AR* to be 1-AR in cases that AR>0.5, thus 0<=AR*<=0.5.

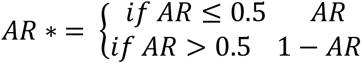

To avoid a noisy signal from a sporadic expression of hSNP, we considered only SNPs that were transcribed in >25% of the cells. We also removed hSNPs that were fully monoallelic to the Ref or the Alt allele (i.e., AR* <0.1). Out of these hSNPs, for each cell we calculate the average of AR*. Cells with an average AR* score of >0.05 show an unexpected biallelic X chromosome expression and were thus considered suspicious as doublets.

### 2.4. Statistical measures for cell doublet identification

For both datasets Mann-Whitney U test was used to determine differences between singlets and doublets according to the BAR values. For Dataset 1 that is based on Smart-seq2 we applied a Gaussian Mixture Model (GMM) that differentiates the groups of singlets from doublets. The GMM was set with two components one seeking the singlets and the other the doublets. The features that were given to the GMM include (i) the Biallelic Ratio (BAR) of each of the cells, and (ii) the number of expressed genes in heterozygous sites in each of the cells.

Dataset 2 (based on 10X Genomics technology) is signified by poor coverage (Supplemental Fig. S4 and Fig. S5), therefore, we included additional features per cell for recovering doublets. The four features that were used are: (i) The number of reads over all heterozygous positions; (ii) The number of expressed genes having heterozygous positions; (iii) The average BAR values; (iv) The fraction of genes defined as biallelic out of all genes expressed in that cell. Each of these features was standardized according to the specific run-individual pair (Kang, et al., 2018). Each of the standardized datasets was trained on its own values. The datasets were split to training and test sets (with the training set covers 75% of the data). We applied Random Forest procedure with the 4 listed features for recording the statistical results. Operating the Random Forest classifier was done with the following parameters: n_estimators=100, random_state=42, min_samples_leaf=sqrt(sample size), min_samples_split=2*sqrt(sample size). Additionally, singlets and doublets were equal weighted by demanding the class_weight to be balanced.

Sensitivity, specificity and accuracy in doublets identification were measured according to the success and failure in detecting simulated and candidate doublets. ROC curve and AUC were calculated for each run-individual pair, for each of the different runs (run_a, run_b and run_c) and for the combined set of all three runs.

### 2.5. Cell separation by gene expression matrix

In this part, we followed the protocol in (Lun, et al., 2016). Count matrix of genes over cells was created for each of the samples using HTSeq (Anders, et al., 2015). The genes to cells matrix was analyzed using SingleCellExperiment Package (Risso, et al., 2018), scater package (McCarthy, et al., 2017) and scran (Lun, et al., 2016). Rtsne package was used to create the t-distributed stochastic neighbor embedding (t-SNE) (Pezzotti, et al., 2017) representation of the 26 first principal components of the PCA of the gene expression profile of each of the run-individual pairs.

## 3 RESULTS

### 3.1 Overview of the BIRD pipeline

In single cells transcriptomics, monoallelic expression of alleles across each of the heterozygous positions is a common phenomenon (Fig.1A). The majority of the hSNPs are monoallelic due to the stochastic nature of expression (Borel, et al., 2015; Reinius and Sandberg, 2015). In doublets if one cell expresses one of the alleles and the other cells the different allele, the result is a shift towards the biallelic expression profile (i.e., 0.1<=AR<0.9). Therefore, a signal with AR centered around 0.5 represents a product of expressing hSNPs derived from both alleles. The key concept underlying BIRD is that doublets can be identified by a signal derived from the shift toward higher biallelic expression ratio (BAR, see Materials and methods, Supplemental Fig. S1). The transformation of each gene and each cell from AR to its average BAR value is illustrated (Fig. 1B, left). The distribution of BAR values from all cells is indicative for the presence of cells that display a substantial biallelic expression and thus are most likely cell doublets (Fig. 1B, right). Testing the performance of BIRD to identify doublets, is based on artificially creating doublets by combining expression profiles from random single cells and testing the potency of statistical methods to correctly identify such in silico simulated doublets (Fig. 1C and Fig. 1D).

**Fig. 1.**
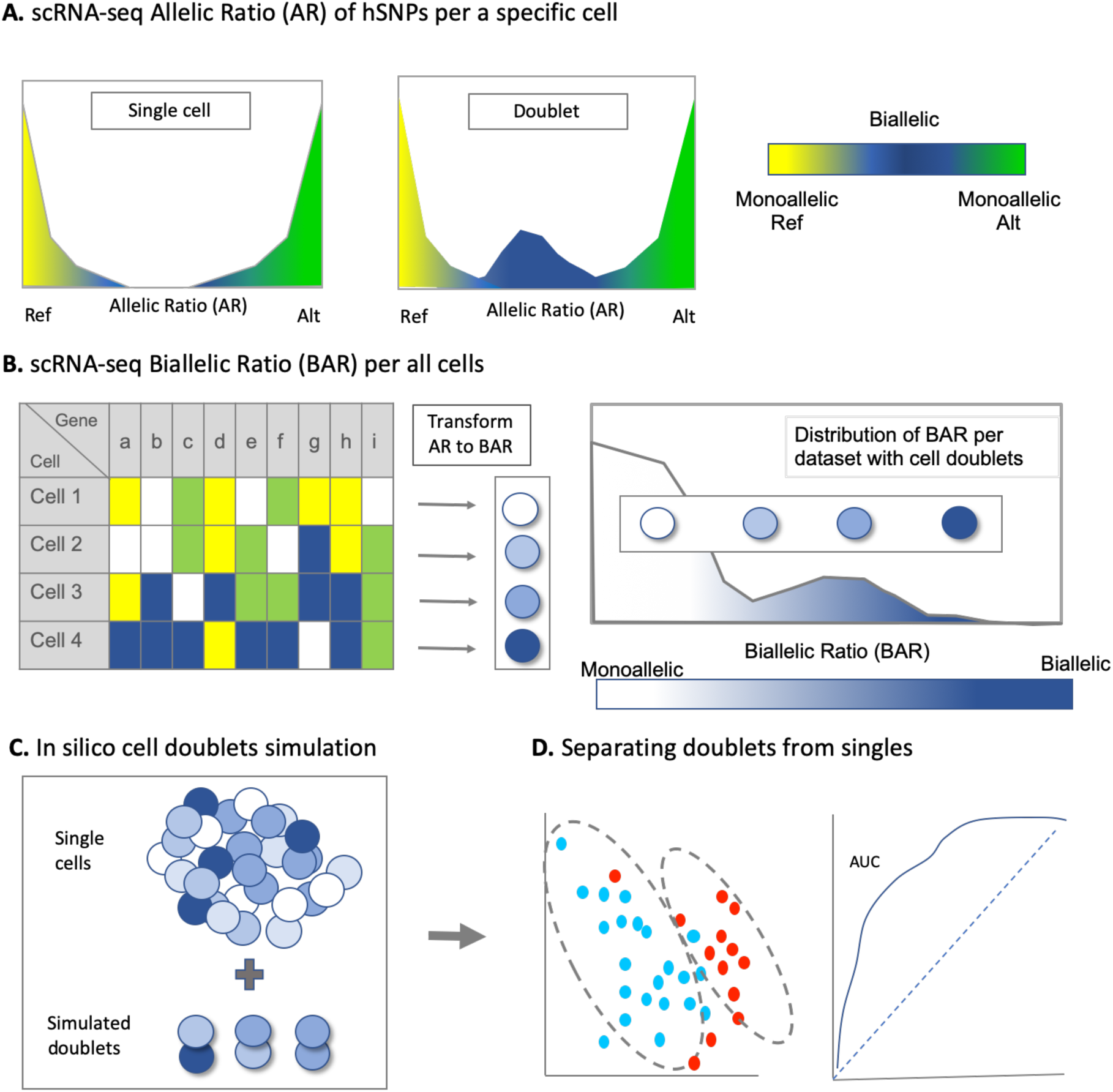
Illustration of the BIRDs scheme for scRNA-seq and dscRNA-seq data. **(A)** Illustrative schemes for the distribution of Allelic Ratio (AR) calculated per each cell. AR values range between 0 to 1, for the reference (Ref, yellow) and Alt (green) alleles, respectively. The blue in the middle corresponds to biallelic expression. (i) For single cells, AR is close to 0 or 1, reflecting an apparent monoallelic expression; (ii) Cell collection with doublets is signified by a shift in AR values to around 0.5 (blue), reflecting biallelic expression pattern. **(B)** On the left, for every gene, in every cell, the AR is estimated. Biallelic Ratio (BAR) for every gene is calculated to create overall BAR distribution for the subjected dataset (see Materials and methods). On the right, the BAR values for scRNA-seq that includes doublets are shown. The BAR is bounded between 0 to 0.5 from monoallelic expression (white) to biallelic (dark blue). **(C)** Simulation of couples of randomly selected single cells is performed to create a dataset composed from both the original cells and simulated cell doublets. **(D)** Machine learning (ML) statistical technique differentiating singlets from doublets. The success of doublet identification is assessed by visualization and standard measures (e.g., AUC).

### 3.2 BAR values for the fibroblast scRNA-seq data

The human primary fibroblast cells (total of 163 single cells) are comprised of two datasets according to the PCR protocol used for creating the sequencing library. The first collection consists of 104 cells that underwent 22 PCR cycles, and the second set consists of 59 cells that underwent 12 PCR cycles. Due to the different PCR protocols, the sequencing depth is different between the two cell collections (Supplementary Fig. S2) and they are thus treated for the BIRD protocol as independent sets. Fig. 2 shows the results of doublet simulations for each of these datasets. The distributions of the means of singlets vs doublets for 100 simulation runs (each with 5% of artificially created doublets) are shown in Fig. 2A and Fig. 2C. Both datasets resulted in a perfect separation with (Mann-Whitney U test statistic=0, and p-value <e-34). The cell values for a single simulation run are also very significant (Fig. 1B and Fig. 1D). The results from the 104 cells (22 PCR cycles) and the 59 cells (12 PCR cycles) show Mann-Whitney U test with a p-value of 4.25e-30 and 1.43e-17, respectively. The results of identifying doublets are data-specific but highly significant for the two cell collections despite the different sequencing depth associated with each.

**Fig. 2.**
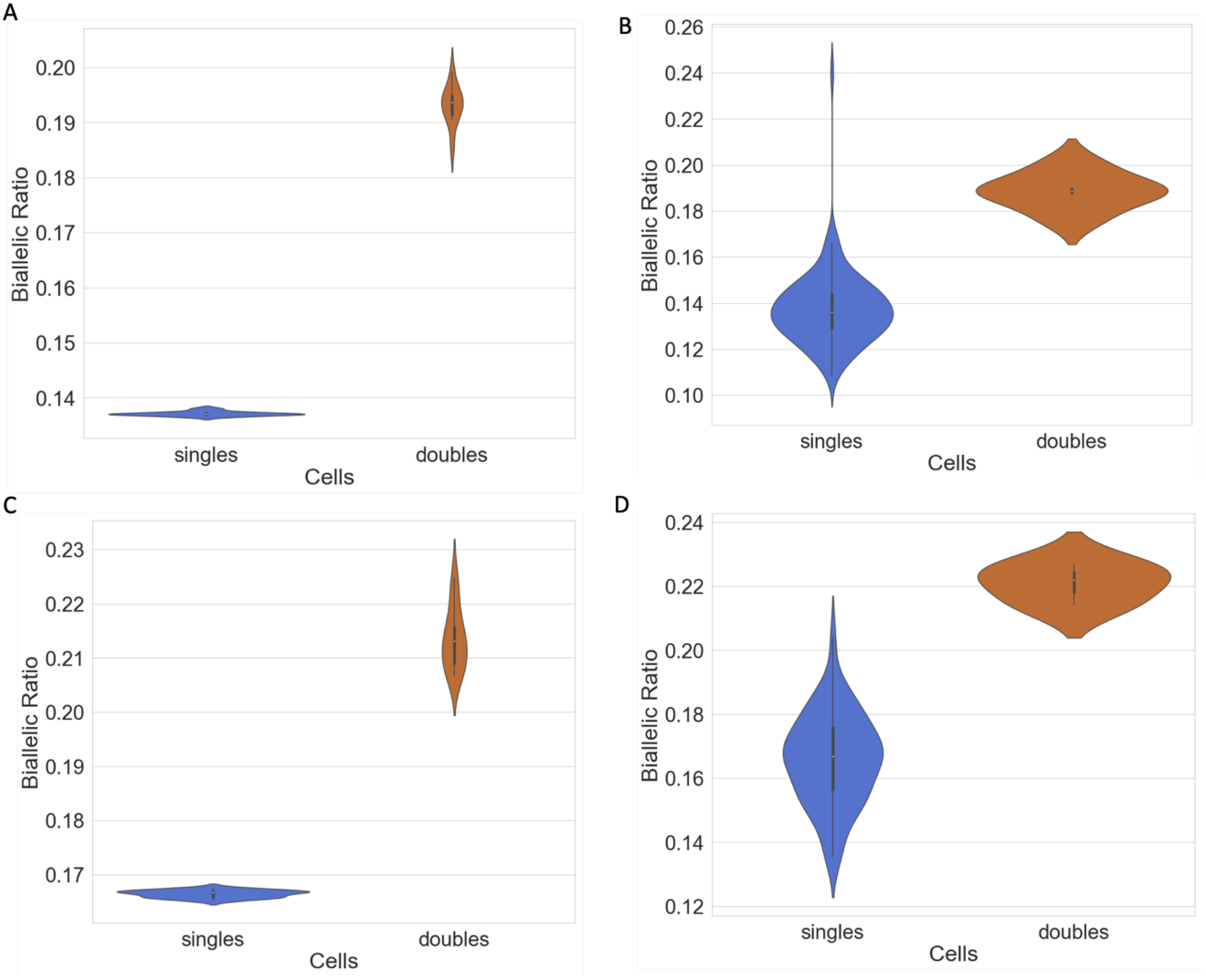
Biallelic ratio (BAR) values for human single cells primary fibroblasts population. Simulations of doublets were done for two datasets of human primary fibroblasts which differ by the number of PCR cycles used prior to sequencing. **(A, B)** 104 cells, 22 PCR cycles and **(C, D)**, 59 cells, 12 PCR cycles. Violin plots of the BAR mean values for all single cells means versus simulated doublets means based on 100 simulations **(A, C)** and cell means of a single simulation **(B, D)**. The applied Mann-Whitney U test results are **(A)** statistic=0, p-value=1.281e-34; **(B)** p-value=4.25e-30; **(C)** statistic=0, p-value=1.281e-34; **(D)** p-value=1.43e-17

### 3.3. Doublets verification based on Chromosome X inactivation expression pattern

The primary fibroblast cells are of a female origin. Thus, in each cell, only one of the two X chromosomes is active (i.e. Xa) while the other is inactivated. The expression patterns for the subset of hSNPs on Chromosome X having substantial evidence are shown (Fig 3A). Most cells (columns) are signified by a single expression pattern that is indicated by Haplotype 1 and Haplotype 2. Only a few cells lean toward biallelic expression pattern over many X chromosome genes. Based on hierarchical clustering of the cells, the cells that are suspicions as doublets are clustered in the leftmost subtree and on the leftmost leaf of the other two subtrees. The distribution of the AR* values for all 163 cells is shown in Fig 3B. AR*=0 means monoallelic X chromosome expression, and the higher the AR*, the higher the biallelic expression is. Applying a natural threshold that separates cells with monoallelic and biallelic patterns (the striped line at AR*=0.05) allows focusing on cells that cross the threshold (8 cells). These cells are marked as cell doublet candidates. Notably, these suspicious 8 cells are also signified by a higher BAR values for the 104 (Mann-Whitney U test p-value=2.94e-4) and 59 (Mann-Whitney U test p-value=0.023) cells as shown in Supplemental Fig. S3.

**Fig. 3.**
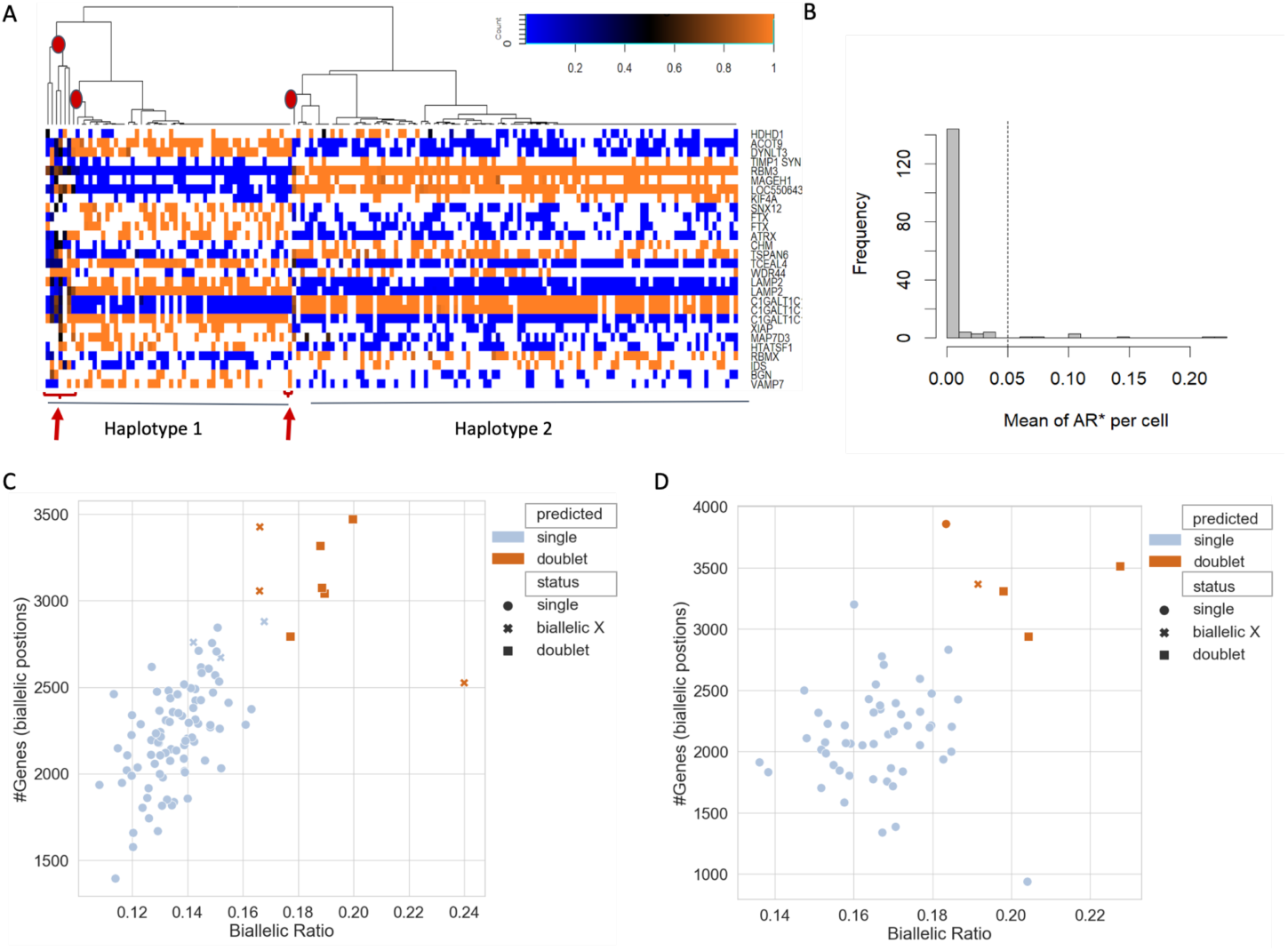
Validation of cell doublets according to X chromosome allelic specific pattern. Dataset is a combined collection of human fibroblasts (163 cells). (**A)** A matrix of AR values of cells (columns) and hSNPs (rows) is shown. Only high support hSNPs are included (see Materials and methods). Gene names are listed according to their chromosomal order (right). The hSNPs are colored from blue (AR=0, Ref allele) to orange (AR=1, Alt allele), with darker colors marking biallelic expression. Hierarchical clustering of the cells indicates two main haplotypic origins (Haplotypes 1 and 2). The arrows and the circles above the branches of the clustering tree indicate cells with strong biallelic expression across the X chromosome. (**B)** Histogram of cells by their AR* values. AR*=0 indicates monoallelic X-chromosome expression, and larger value marks a higher biallelic expression level. The dashed line is a natural threshold separating X-inactivated monoallelic from biallelically expressed hSNPs. **C, D** Scatter plots show the success of detecting singlets from doublets following a single simulated set for the 104 cells **(C)** and 59 cells **(D)**. Symbols correspond to singlets as circles and doublets as squares. A cell is marked by x if it was validated as a doublet on the background of the X-inactivated status (as in **A, B**). Dark orange marks cells that were identified by the GMM classifier (see Materials and methods) as doublets.

### 3.4. Unsupervised identification of doublets for the fibroblast scRNA-seq data

Gaussian Mixture Model (GMM) was used to separate singles from doublets. For the means of the 100 simulations, the separation between the singles and the doublets means reached 100% accuracy. For an illustrative of a single simulation run the mean BAR (x axis) of each cell is plotted with the number of genes that are expressed in biallelic positions (y-axis, Fig. 3C, and Fig. 3D). The scatter-plots symbols represent cells that are singlets, candidate cell doublets according to Chromosome X biallelic expression, and artificial doublets that are created by in silico simulations for the two fibroblast cell collections (104 and 59 cells based on PCR protocol for 22 and 12 cycles, respectively). Cells that were predicted as doublets by the GMM (whether true or false) are shown (dark orange). It is evident that most doublets and the Chromosome X candidate doublets have relatively high BAR values and are classified as doublets. For the 104 single cells dataset, all simulated doublets (total 5) were identified (100%), and 3/6 (50%) of the candidates by X biallelic expression were identified. For the 59 cells, all 3 simulated doublets were identified (100%) and 50% of the cell candidate doublets (one out of two) according to the Chromosome X biallelic expression were correctly identified.

### 3.5 BAR values for the peripheral blood mononuclear cells (PBMCs) dscRNA-seq data

The 13,364 peripheral blood mononuclear cells (PBMCs) originate from 16 datasets that account for a pair of a run and an individual. When compared to the fibroblast cell collections (Dataset 1, see Materials and methods), the dscRNA-seq is characterized by a much lower coverage (Supplemental Fig. S4 and S5). Specifically, the number of informative genes is >2000 for the fibroblasts and only about 25 on average for the PBMCs (Supplemental Fig. S2, Fig. S4, and Fig. S5).

We simulated as in the fibroblasts, for each of the datasets, 10% of the singles to be paired into in silico doublets (overall 5% of each dataset would be doublets). BIRD was run on each of the 16 run-individual pairs and their in silico simulated doublets. The results from activating the BIRD process for an individual representative (run_b, individual 1493, denoted b_1493, 766 cells) are shown in Fig. 4. (and Supplemental Fig S4).

**Fig. 4.**
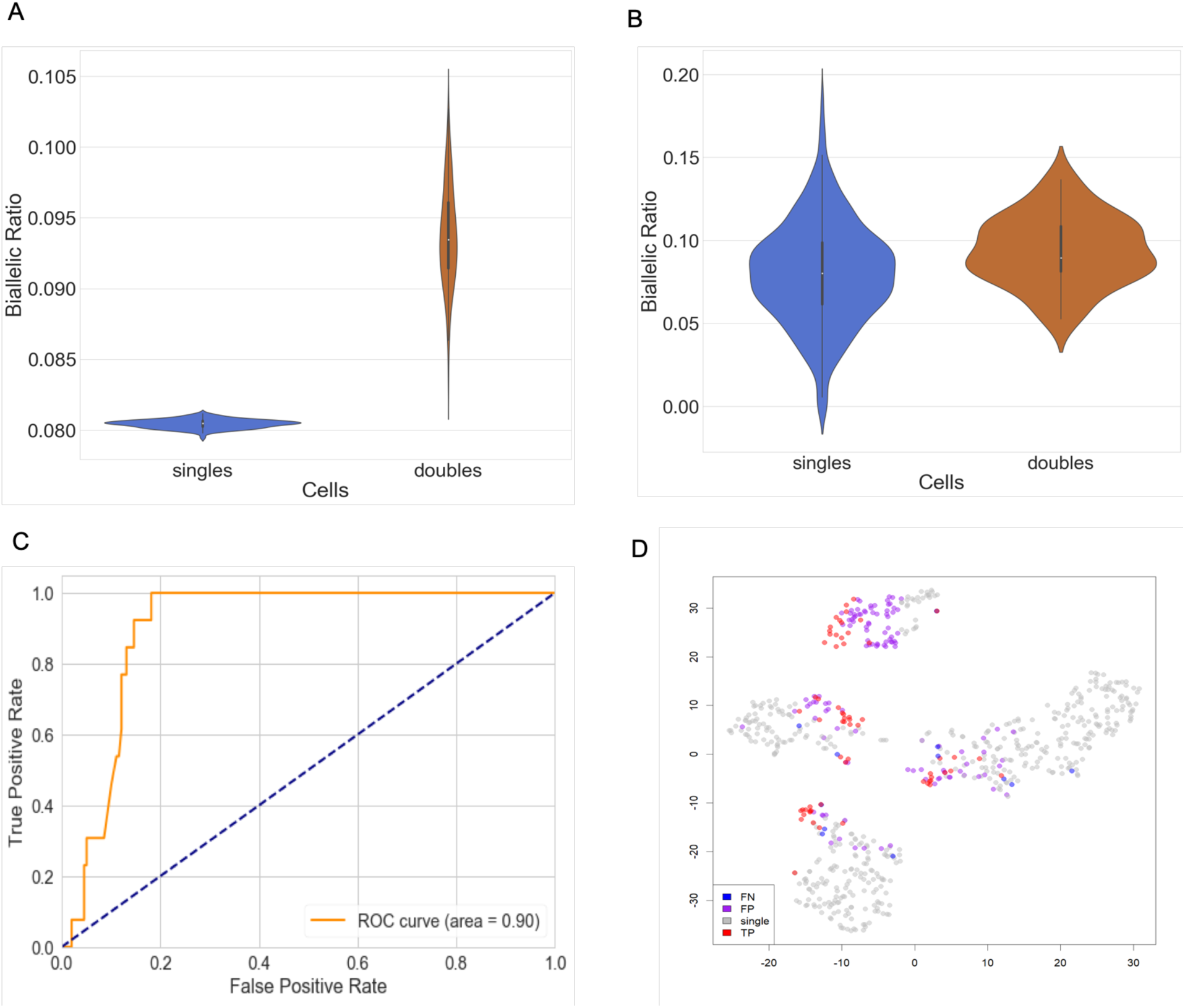
Data used is dscRNA-seq from a run-individual pair marked as b_1493 based on Dataset 2 (See Materials and methods). Violin plots of the mean of BAR values for single cells and the in silico formed doublets. The mean BAR values shown were tested by Mann-Whitney U test for **(A)** 100 simulations (statistic=0, p-value <e-34), and **(B)** a single simulation set (p-value=1.22e-03). (**C**) A receiver operating characteristic (ROC) curve is shown based on a Random Forest (RF) model fitting on a simulated dataset of singlets and doublets. (**D)** t-distributed stochastic neighbor embedding (t-SNE) classification on PCA reduced data of cell expression (see Materials and methods). Each dot represents a single cell or a simulated doublet. Singlets (gray) corresponds to cells that are singles and were predicted by the model as singles. True positives (TP, red) correspond to cells that are simulated as doublets and correctly predicted as such. False Negatives (FN, blue) are simulated doublets that were missed by the model. False Positives (FP, purple) are misclassified by the model as doublets. Note that most of the identified doublets cluster in a cloud of cells that are likely to represent mixtures of cell types.

Fig 4A shows the violin plots of the mean of the BAR values for 100 simulations for singlets when compared to cell doublets. For the means of the singles and the doublets, the separation is maximal with Mann-Whitney U test yields a statistic=0, and a p-value <e-34. For a single run of the simulation (Fig 4B), the trend of the doublets being more biallelic is kept with a separation of the Mann-Whitney U test yielding a p-value of 1.22e-03. Note that using BAR values alone is insufficient to distinguish between singles and doublets due to the high intrinsic noise in the data originated by the 10x Genomics protocol.

### 3.6 supervised identification of doublets in the PMBC dataset

Including additional features and applying a supervised Random Forest machine learning protocol (see Materials and methods), we reached a perfect separation with 100% accuracy when mean values of singlets and mean values of simulated doublets are compared for 100 simulation runs. For a single simulation run, we show the results of the Receiver operating characteristics (ROC) curve for the unseen, disjoint test. The Area Under the Curve (AUC) of the ROC curve equals 0.9 (Fig 4C). As each run starts with a randomized set of simulated doublets, we report the average AUC for 10 independent such runs (mean 0.88, s.d. 0.04). We exploit the expression profile for the cell collection of b_1493 sample to create a t-SNE representation (Fig 4D). The cells are color coded according to the prediction results. Note that most identified doublets are positioned at the border of the expression clusters, but eventually other predicted doublets are fully embedded within an expression cluster. Recall that the expression profile information was not used by BIRD protocol for the separation of the prediction.

Similar to the analysis performed for a single dataset (b_1493) we repeated the analysis for all 16 combinations of runs and individuals. The AUC values for the different datasets of the run-individual pairs (see Supplementary Table S1) are shown in Fig 5A. We tested whether the performance (as indicated by the AUC) is a mere reflection of the number of cells. However, it is evident that the success in identifying doublets and the number of cells that are associated with each dataset are not correlated (Supplemental Fig. S6A). The sensitivity (i.e. TP/(TP+FN)) for each of the 16 datasets is shown in Fig 5B and Supplementary Table S1. Note that the values in Supplementary Table S1 for dataset b_1493 are slightly different from results in Fig. 4 due to the component of randomness in the algorithm. The doublets rate in the sample (including the simulated cells) is shown in Supplemental Fig. S6B. We created a unified ROC curve per each of the three runs and determined the AUC associated with each run and the whole dataset (Fig 5C).

**Fig. 5.**
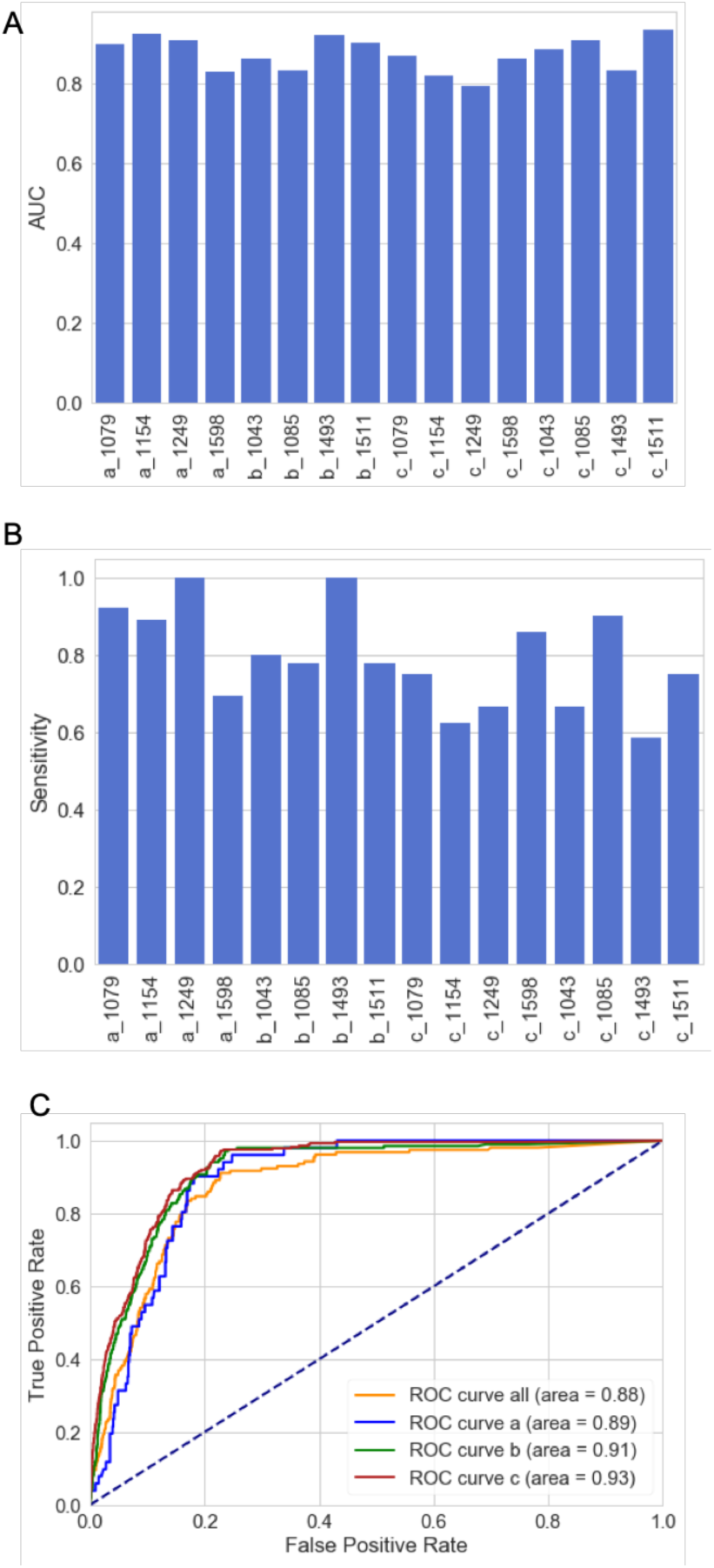
Success in identifying cell doublets from multiplex 10X Genomics experiment covering 13,364 single cells. (**A**) The Area under the curve (AUC) for the test sets of each of run-individual pairs. Runs refer to run_a, run_b that consist of 4 different individuals each, and run_c that combines all 8 individuals. (**B)** The sensitivity achieved for the test set for each of the tested individuals. **(C)** ROC curve is shown based on a Random Forest (RF) model fitting on a simulated dataset of singles and doublets for all the cells (marked all) and for each of the separated runs (a, b and c).

The t-SNE cell expression representations are colored according to the BIRD prediction and the Random Forest protocol, for each of the 16 datasets are shown in Supplemental Fig S7 for run_a and run_b (4 t-SNE representations per each run) and Supplemental Fig S8 for run_c (all individuals, 8 t-SNE representations). In all instances, the t-SNE representation shows that accurate predictions (TP) tend to cluster together with cells that are marked as false positives (FP). The estimate for the fraction of doublets from the mixture of two individuals is ∼5% (Kang, et al., 2018). Therefore, we expect many of the cells that are marked as FP to be doublets that are naturally present in the original data.

## 4 DISCUSSION

Collecting data of single-cell transcriptomes had exposed a new dimension of cell variability. This technology had a direct impact on a wide range of biological questions across all domains of life (Stegle, et al., 2015). Some of these questions are sensitive to the faulty annotation of singlets as doublets or vice versa. While the presence of unrecognized cell doublets from the same cell type will not influence the misinterpretation for new cell types (Usoskin, et al., 2015; Villani, et al., 2017; Zeisel, et al., 2015), it might jeopardize interpretation concerning transcription regulation including transcriptional bursting kinetics (Larsson, et al., 2019), Chromosome X-inactivation phenomenon (Garieri, et al., 2018; Tukiainen, et al., 2017), escaping from it (Wainer-Katsir and Linial, 2019) and more.

We describe BIRD as a computational/statistical method that enables the identification of cell doublets from scRNA-seq data. The method complements other methods that rely on detailed cell mixing and cell-type expression profiles. BIRD method takes advantage of the BAM files generated for each scRNA-seq. The allelic specific expression that is extracted from the BAM files is often unused. This is since, routinely, the post-sequencing analysis starts with a cell to gene matrix representation thus discards the allelic information. BIRD takes advantage of this transparent feature for identifying doublets.

Recent biochemical-based methodologies for dscRNA-seq present their potency toward the task of doublet identification. These methods are based on adding a pre-sequencing biochemical modification step for barcoding cells. Such tagging procedures exploit antibodies to common cell surface antigens (cell hashing) (Stoeckius, et al., 2018). The antibody-based method applies to cells that carry the relevant antigens. A new method (MULTI-seq) (McGinnis, et al., 2019) successfully uses lipid modification step for cell indexing. It was shown to be eligible for solid tissues and frozen cells (McGinnis, et al., 2019). While there are many advantages for tagging cells before cell lysis and sequencing, an additional step in the experimental design can lead to batch effect and other technical and experimental biases. In contrast, the computational method is generic, yet data sensitive. BIRD shows no preference to the identity of the expressed genes, to the specific cell type or any of the cell extraction protocol.

We illustrate the high performance of BIRD mostly on in silico simulated doublets. Other studies estimated the rates of doublets by artificial mixing of cells of multiple types of cells from different organisms, cell types or individuals (Kang, et al., 2018; McGinnis, et al., 2019; Zheng, et al., 2017). However, when a solid tissue is treated to produce a collection of single cells, the protocol must overcome the adhesion forces between cells, extracellular matrix cohesion and more. Additionally, some cells tend to aggregate and clump following their isolation. All these technical issues may lead to an increasing number of doublets from neighboring cells with identical genetic background and expression profile. Therefore, current estimates for doublet contamination based on peripheral blood samples may be misleading. We anticipate that the number of reported doublets of cell mixtures from solid tissue is underestimated and can now be estimated using BIRD.

There are several limitations of BIRD protocol that need to be addressed: (i) The method is dependent on pre-knowledge of the individual genomics for assigning hSNPs from the sequenced scRNA-seq. With the fast accumulation of whole-genome and exome sequencing in humans and other model animals, we anticipate it will not be a limiting factor in the near future. (ii) The assessment of doublets using the notion of Chromosome X inactivation is only valid for cells of female origin. Furthermore, for 50% of the cases, cells can be mixed without providing a biallelic signature (i.e. a mixture of the same Xa haplotype). (iii) While BIRD protocol ignores the gene expression profile, a scenario in which the expression profiles of cell mixtures do not overlap with each other can occur. This will result in cell doublets that do not contribute different alleles of the same genes and thus will not increase the BAR values. In such cases, BIRD protocol lacks the power to identify doublets. In this case, the use of other doublet cell identification is advisable (e.g. (McGinnis, et al., 2019; Wolock, et al., 2019)). (iv) The method relies on the dominant properties of stochasticity in the allelic expression of cells. Datasets that are far less stochastic might display a higher biallelic signal. Under such conditions, the ability to detect doublets is masked. The monoallelic fraction in single cells is a variable property of the experiment (Kim, et al., 2015) with mouse cells showing a lower degree of allelic specific expression relative to humans (Deng, et al., 2014; Tang, et al., 2011). Indeed, testing the scRNA-seq from F1 mice strains (Larsson, et al., 2019), confirms that the BAR value distribution is consistent with an intrinsic biallelic signature (not shown). In such cases, there is a need to employ BIRD only on genes that exhibit a more stochastic property and are signified by a monoallelic expression.

Overall, we described two types of datasets that use the BAR feature for discriminating singlets from doublets. The overall coverage of hSNPs and sequence depth are drastically different (Supplementary Fig. S2 Fig. S4 and Fig. S5) among the two analyzed datasets. The unsupervised GMM tool was sufficient in separating singles from doublets in a dataset of high hSNP coverage (based on Smart-seq2 technology, with a full transcript sequencing). The other dataset (dataset 2, 10X Genomics) yields shallow coverage which is restricted to the 3’ tail of the transcripts and was therefore trained using Random Forest. Despite the described coverage and difference in the sequencing protocols (3’ based versus a full-length transcript), the mean values of the simulated doublets were 100% identifiable for both datasets indicating a higher BAR values for doublets compared to singlets. Even in the cases of a single run, a trend of higher BAR values for doublets was observed and therefore used to successfully separate the simulated doublets from the singlets in both datasets.

In summary, BIRD protocol is a generic method that is indifferent to the composition of the cells in the samples of to the nature of the genes that are expressed. It is on the other hand strongly dependent on the stochasticity of the system for identifying doublets according to the deviation from the default signal of monoallelic expression that dominate single cells. We applied BIRD on datasets of different coverage, scale, and accuracy. BIRD uses a data-driven protocol and is applicable in all instances where in addition to the scRNA-seq / dscRNA-seq data genomic heterologous sites are available.

## ACKNOWLEDGEMENTS

We thank Nadav Brandes for excellent discussion, the Linial lab for useful comments, Liran Carmel (HUJI) for a statistical support and Oren Ram (HUJI) for sharing the challenges of single cell genomics. KWK is a recipient of the Ariane de Rothschild Scholarship. This study used best practice protocols and education and training materials presented by EU-Elixir.

## SUPPLEMENTAL FIGURE LEGENDS

**Table S1.** BIRD performance of 16 run-individual pairs covers 13,364 cells by Random Forest prediction.

| **sample_**<br>**name** | **#Cells per**<br>**sample** | **Accuracy** | **Sensitivity** | **Specificity** | **Precision** | **Doublet**<br>**rate<sup>a</sup>** | **AUC** |
| --- | --- | --- | --- | --- | --- | --- | --- |
| a_1079 | 886 | 0.79 | 0.92 | 0.78 | 0.20 | 0.26 | 0.90 |
| a_1154 | 999 | 0.83 | 0.89 | 0.82 | 0.15 | 0.20 | 0.92 |
| a_1249 | 810 | 0.84 | 1.00 | 0.83 | 0.24 | 0.22 | 0.91 |
| a_1598 | 800 | 0.86 | 0.69 | 0.87 | 0.26 | 0.16 | 0.83 |
| b_1043 | 1057 | 0.76 | 0.80 | 0.76 | 0.11 | 0.26 | 0.86 |
| b_1085 | 1038 | 0.81 | 0.78 | 0.82 | 0.23 | 0.22 | 0.83 |
| b_1493 | 766 | 0.88 | 1.00 | 0.87 | 0.26 | 0.17 | 0.92 |
| b_1511 | 1227 | 0.88 | 0.78 | 0.88 | 0.29 | 0.15 | 0.90 |
| c_1079 | 730 | 0.86 | 0.75 | 0.87 | 0.27 | 0.17 | 0.87 |
| c_1154 | 757 | 0.76 | 0.63 | 0.76 | 0.10 | 0.25 | 0.82 |
| c_1249 | 648 | 0.87 | 0.67 | 0.88 | 0.24 | 0.15 | 0.79 |
| c_1598 | 648 | 0.84 | 0.86 | 0.83 | 0.18 | 0.19 | 0.86 |
| c_1043 | 771 | 0.85 | 0.67 | 0.86 | 0.13 | 0.16 | 0.88 |
| c_1085 | 765 | 0.84 | 0.90 | 0.83 | 0.22 | 0.20 | 0.91 |
| c_1493 | 619 | 0.82 | 0.58 | 0.84 | 0.23 | 0.19 | 0.83 |
| c_1511 | 843 | 0.87 | 0.75 | 0.88 | 0.10 | 0.14 | 0.93 |
| **Total_cells** | **13364** |  |  |  |  |  |  |
| **Average** | **835.3** | **0.83** | **0.79** | **0.84** | **0.20** | **0.19** | **0.87** |
| **s.d.** | 167.9 | 0.038 | 0.128 | 0.041 | 0.064 | 0.040 | 0.043 |

<sup>a</sup>Double rate applies to cells identified as doublets (including simulated doublets) out of the total cells per sample.

**Fig. S1.**
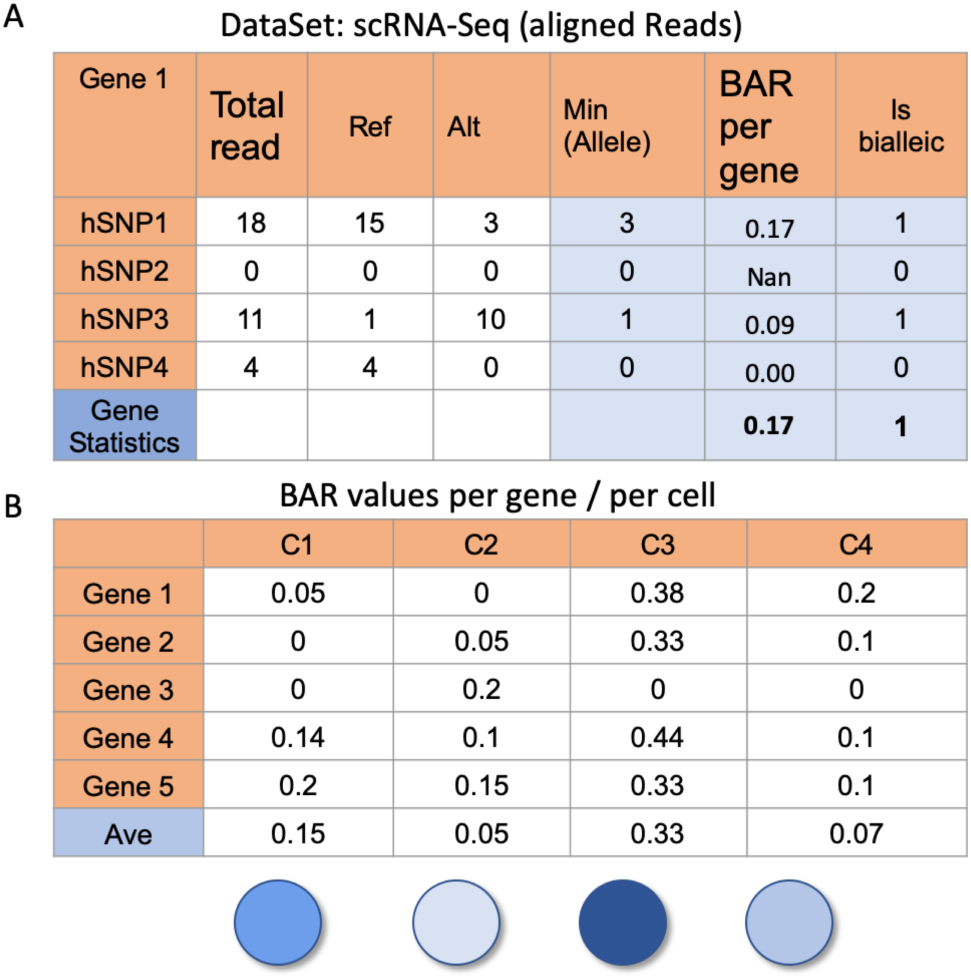
The numeric transition from reads per hSNP to BAR of cells. (A) In each cell for each gene for each expressed SNP, the BAR is determined. First, the number of Alternative (Alt), Reference (Ref) and total reads are counted for each position. The algorithm takes the BAR of an hSNP to be the ratio between the reads of the minimal expressing allele divided by the total reads of that position. Then for each gene, the hSNP with the maximal value of BAR is chosen as a representative hSNP BAR (as indicated by the bottom row). Additionally, each hSNP is recorded as biallelic or not. And one representation of a biallelic hSNP determines the gene as biallelic (rightmost column). (B) For each gene, in each cell, the BAR is recorded. For a specific cell, it’s BAR distribution will be according to the BAR of its genes. The average of the genes BAR will define the cells BAR.

**Fig S2.**
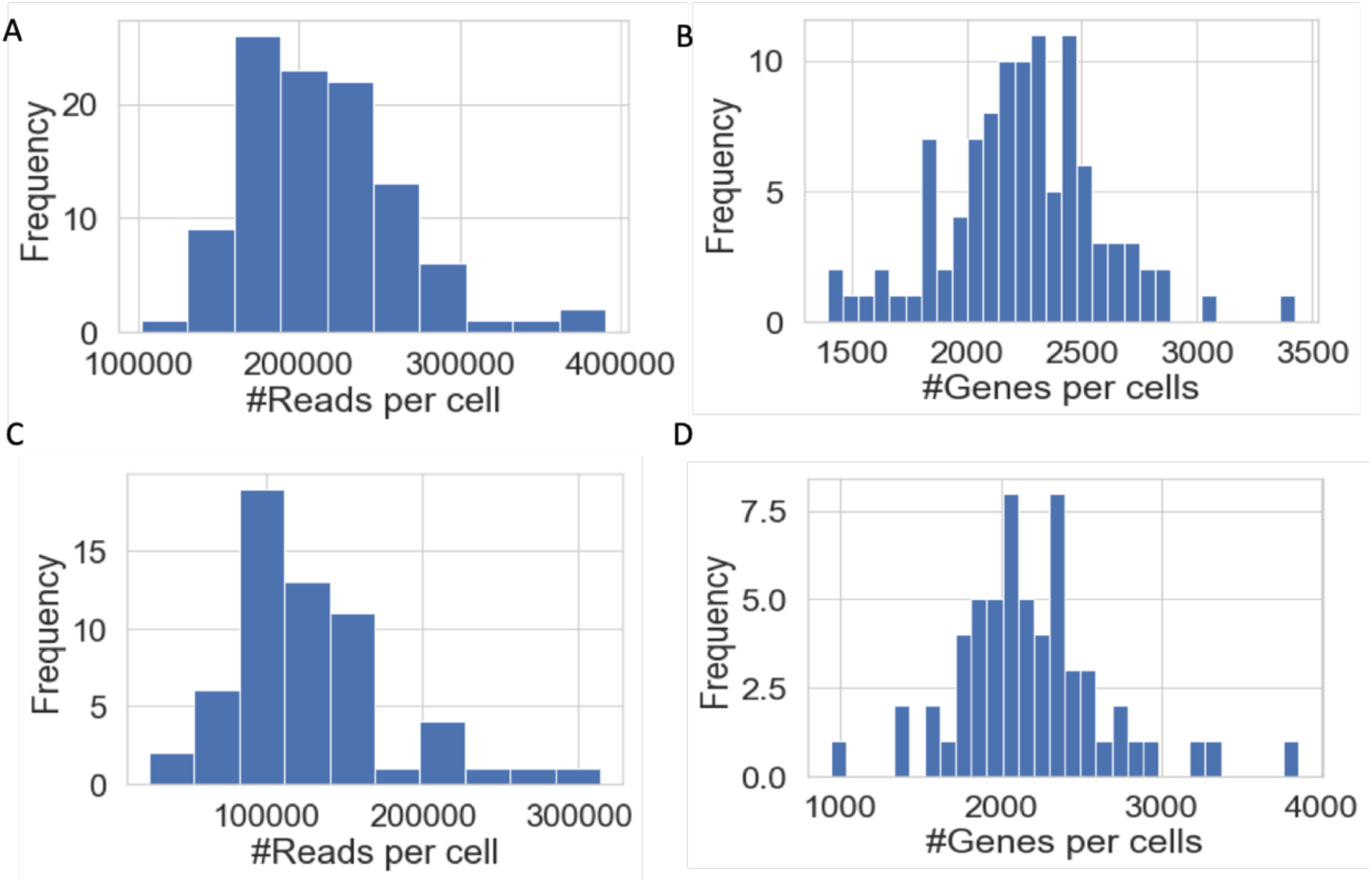
Coverage of fibroblast dataset. **(A, B)** 22 PCR cycles dataset with 104 cells. **(A)** Histogram of the number of reads per cell in heterozygous positions. **(B)** Histogram of the number of genes per cell in heterozygous positions. **(C, D)** 12 PCR cycles dataset with 59 cells. **(C)** Histogram of the number of reads per cell in heterozygous positions. **(D)** Histogram of the number of genes per cell in heterozygous positions.

**Fig S3.**
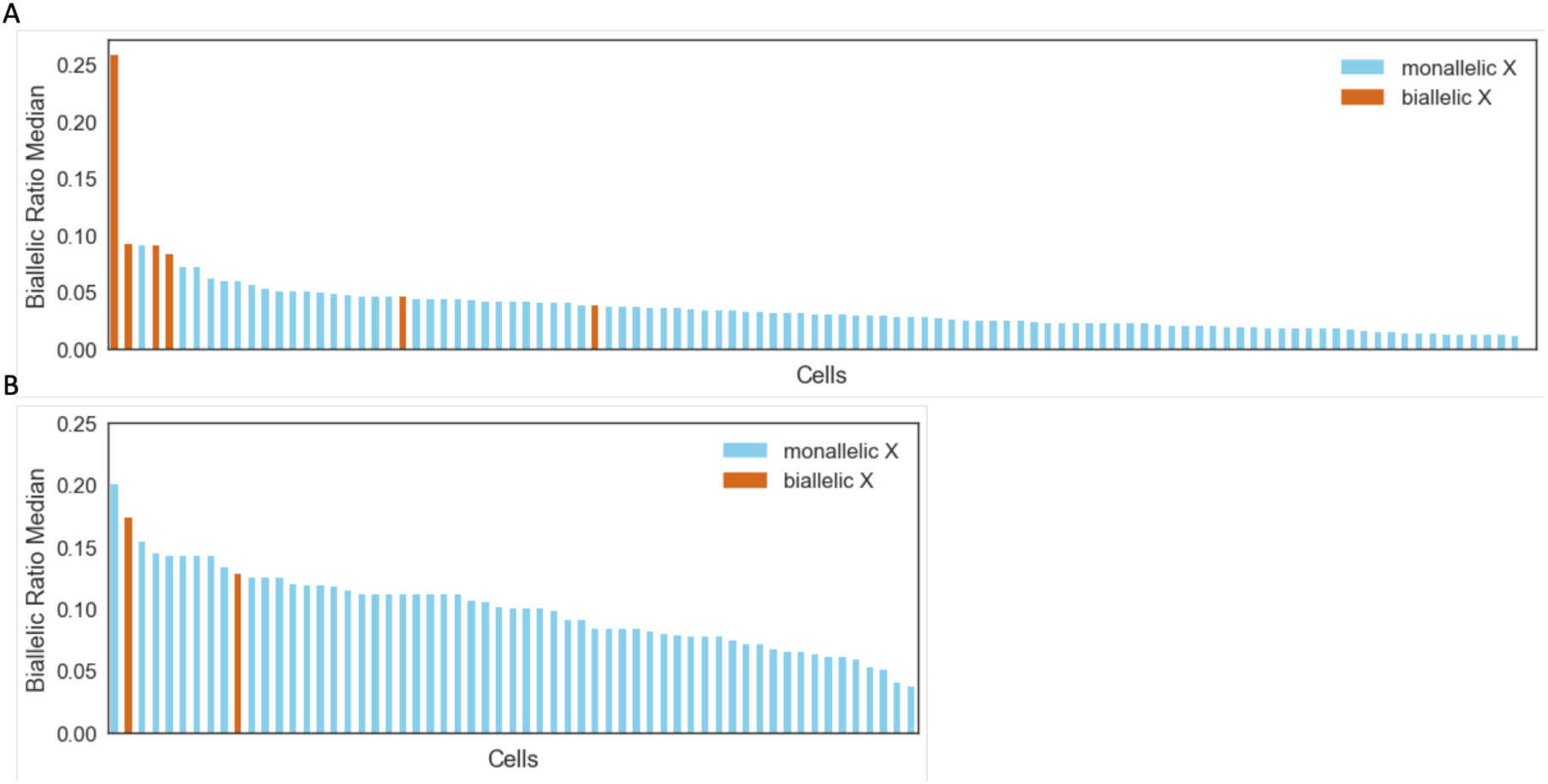
BAR median values of fibroblast dataset. (A, B) Cells are sorted according to their Biallelic Ratio Median. Blue indicates a single according to the X inactivation verification step. Brown cells are doublets according to the X inactivation verification step. (A) Cells from 22 PCR cycles 104 dataset. (B) Cells from 12 PCR cycles 59 cells dataset.

**Fig S4.**
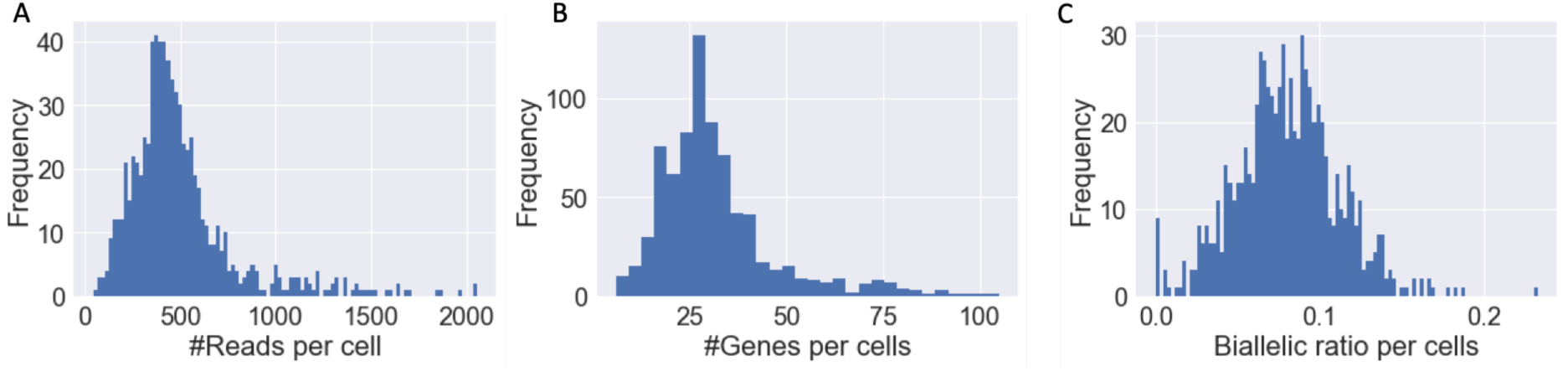
Coverage of a representative PMBC sample of b_1493 (766 cells). (A) Histogram of the number of reads per cell in heterozygous positions. (B) Histogram of the number of genes per cell in heterozygous positions. (C) Histogram of the BAR of the cells.

**Fig S5.**
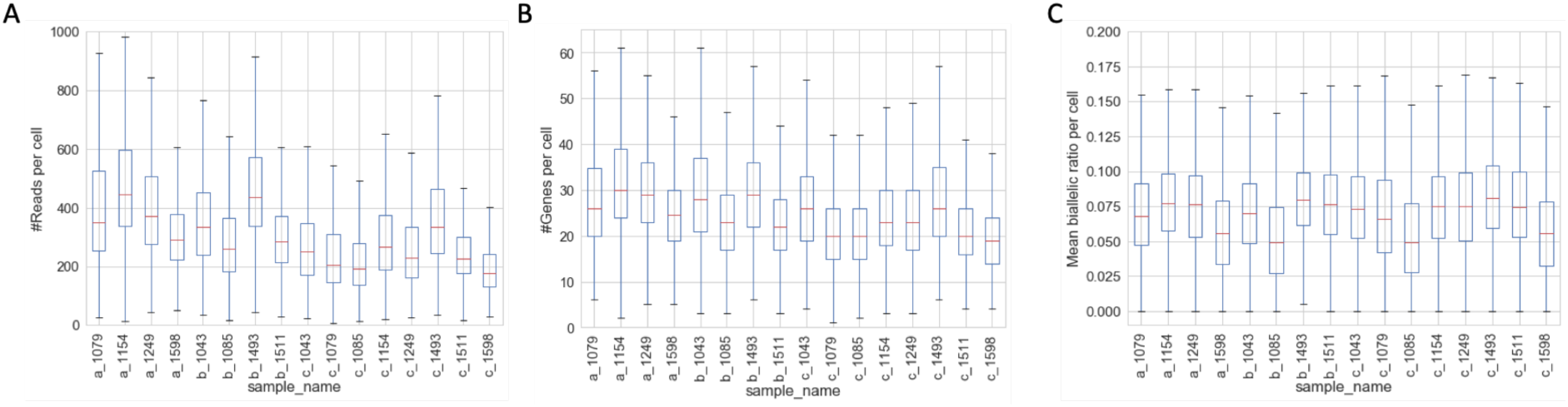
The coverage of all PMBC run-individual pairs. The boxplots show the distribution as in Fig. S4 for each of the run-individual pairs. (A) The number of reads per cell in heterozygous positions. (B) Number of genes per cell in heterozygous positions. (C) The BAR of the cells.

**Fig S6.**
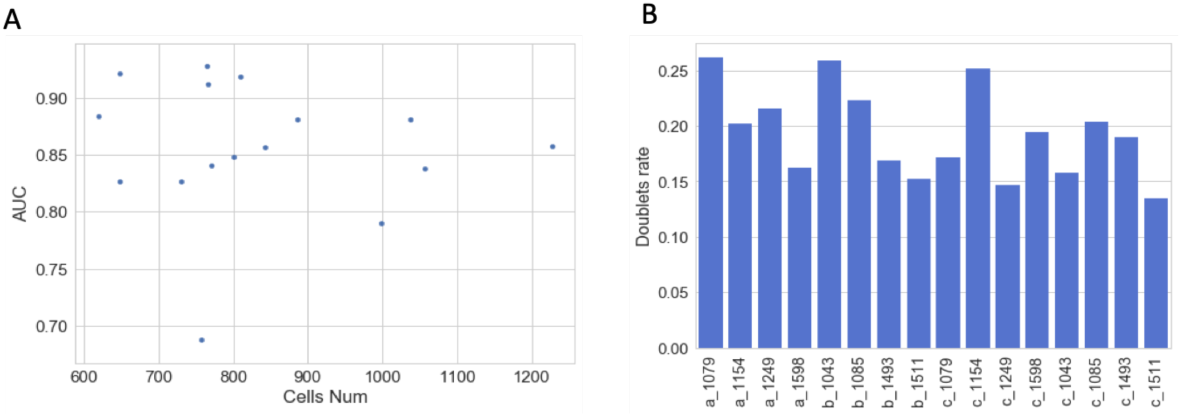
Statistic measures of the Random Forest model results for the single and simulated doublets of the PMBC data. (A) The AUC for the different run-individual pairs (x-axis) by the number of cells per this sample (y-axis). (B) The doublets rate of the test set (including simulated doublets).

**Fig S7.**
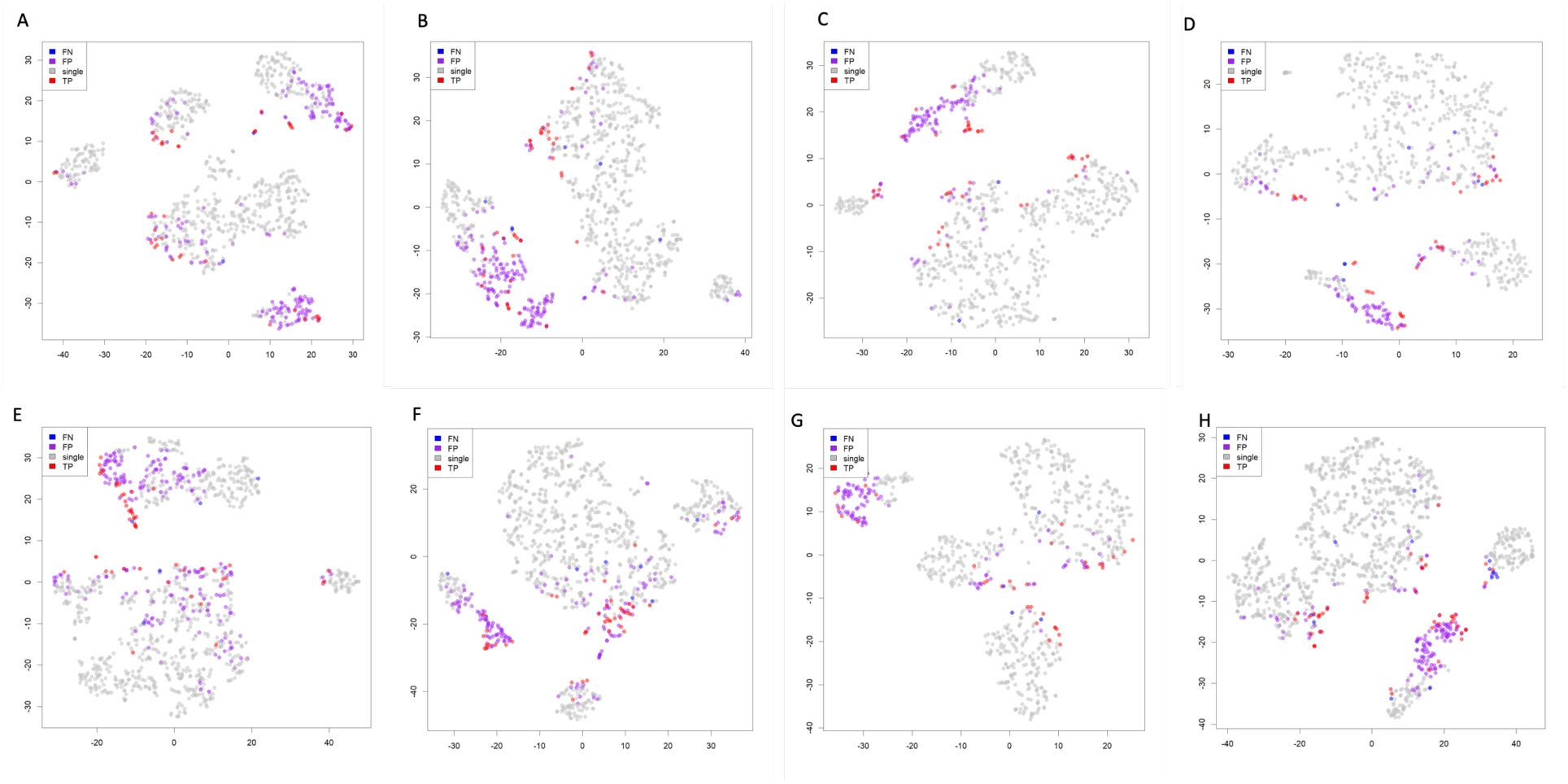
Representation of t-distributed stochastic neighbor embedding (t-SNE) classification on PCA reduced data of cell expression for each of the run-individual paired samples in run_a and run_b (4 individuals per each run). Each dot represents a single cell or a simulated doublet. Singlets (gray) corresponds to cells that are singles and were predicted by the model as singles. True positives (TP, red) correspond to cells that are simulated as doublets and correctly predicted as such. False Negatives (FN, blue) are simulated doublets that were missed by the model. False Positives (FP, purple) are misclassified by the model as doublets. For the individual-run pairs of (A) a_1079, (B) a_1154, (C) a_1249, (D) a_1598, (E) b_1043, (F) b_1085, (G) b_1493, and (H) b_1511.

**Fig S8.**
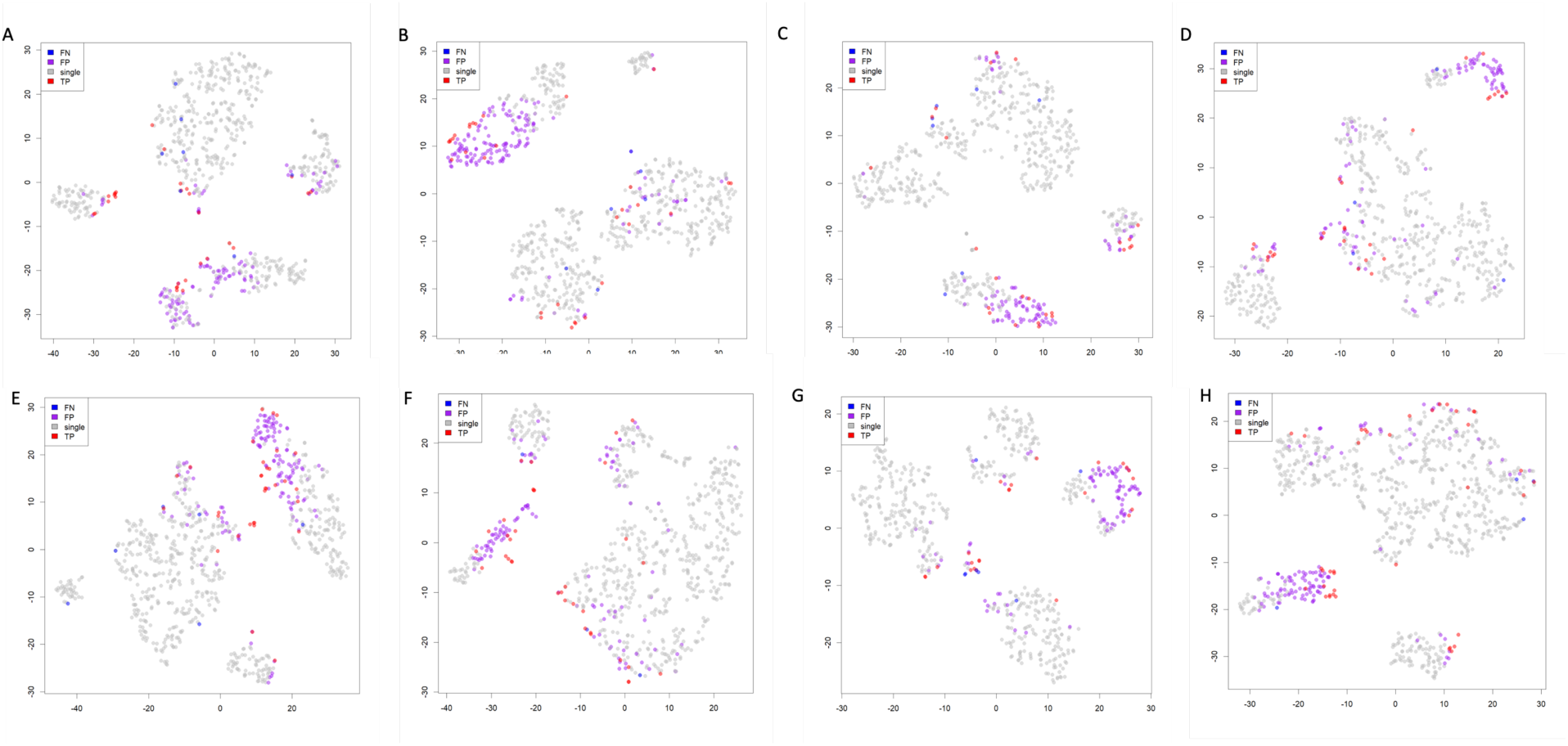
Representation of t-distributed stochastic neighbor embedding (t-SNE) classification on PCA reduced data of cell expression for each of the 8 run-individual paired samples in run_c (8 individuals). Each dot represents a single cell or a simulated doublet. Singlets (gray) corresponds to cells that are singles and were predicted by the model as singles. True positives (TP, red) correspond to cells that are simulated as doublets and correctly predicted as such. False Negatives (FN, blue) are simulated doublets that were missed by the model. False Positives (FP, purple) are misclassified by the model as doublets. For the individual-run pairs of (A) a_1079, (B) a_1154, (C) a_1249, (D) a_1598, (E) b_1043, (F) b_1085, (G) b_1493, and (H) b_1511.

## REFERENCES

Anders, S., Pyl, P.T. and Huber, W. HTSeq--a Python framework to work with high-throughput sequencing data. Bioinformatics 2015;31(2):166–169.

Bacher, R. and Kendziorski, C. Design and computational analysis of single-cell RNA-sequencing experiments. Genome Biol 2016;17:63.

Bolger, A.M., Lohse, M. and Usadel, B. Trimmomatic: a flexible trimmer for Illumina sequence data. Bioinformatics 2014;30(15):2114–2120.

Borel, C., et al. Biased allelic expression in human primary fibroblast single cells. Am J Hum Genet 2015;96(1):70–80.

Buettner, F., et al. Computational analysis of cell-to-cell heterogeneity in single-cell RNA-sequencing data reveals hidden subpopulations of cells. Nat Biotechnol 2015;33(2):155–160.

Castel, S.E., et al. Tools and best practices for data processing in allelic expression analysis. Genome Biol 2015;16:195.

Chen, G., Ning, B. and Shi, T. Single-Cell RNA-Seq Technologies and Related Computational Data Analysis. Front Genet 2019;10:317.

Deng, Q., et al. Single-cell RNA-seq reveals dynamic, random monoallelic gene expression in mammalian cells. Science 2014;343(6167):193–196.

Dobin, A. and Gingeras, T.R. Mapping RNA-seq Reads with STAR. Curr Protoc Bioinformatics 2015;51:11 14 11–19.

Fan, H.C., Fu, G.K. and Fodor, S.P. Expression profiling. Combinatorial labeling of single cells for gene expression cytometry. Science 2015;347(6222):1258367.

Garieri, M., et al. Extensive cellular heterogeneity of X inactivation revealed by single-cell allele-specific expression in human fibroblasts. Proc Natl Acad Sci U S A 2018;115(51):13015–13020.

Haque, A., et al. A practical guide to single-cell RNA-sequencing for biomedical research and clinical applications. Genome Med 2017;9(1):75.

Hashimshony, T., et al. CEL-Seq2: sensitive highly-multiplexed single-cell RNA-Seq. Genome Biol 2016;17:77.

Ilicic, T., et al. Classification of low quality cells from single-cell RNA-seq data. Genome Biol 2016;17:29.

Jiang, Y., Zhang, N.R. and Li, M. SCALE: modeling allele-specific gene expression by single-cell RNA sequencing. Genome Biol 2017;18(1):74.

Kang, H.M., et al. Multiplexed droplet single-cell RNA-sequencing using natural genetic variation. Nat Biotechnol 2018;36(1):89–94.

Kim, J.K., et al. Characterizing noise structure in single-cell RNA-seq distinguishes genuine from technical stochastic allelic expression. Nat Commun 2015;6:8687.

Klein, A.M., et al. Droplet barcoding for single-cell transcriptomics applied to embryonic stem cells. Cell 2015;161(5):1187–1201.

Kolodziejczyk, A.A., et al. The technology and biology of single-cell RNA sequencing. Mol Cell 2015;58(4):610–620.

Lan, F., et al. Single-cell genome sequencing at ultra-high-throughput with microfluidic droplet barcoding. Nat Biotechnol 2017;35(7):640–646.

Larsson, A.J.M., et al. Genomic encoding of transcriptional burst kinetics. Nature 2019;565(7738):251–254.

Lun, A.T., Bach, K. and Marioni, J.C. Pooling across cells to normalize single-cell RNA sequencing data with many zero counts. Genome Biol 2016;17:75.

Lun, A.T., McCarthy, D.J. and Marioni, J.C. A step-by-step workflow for low-level analysis of single-cell RNA-seq data with Bioconductor. F1000Res 2016;5:2122.

Macaulay, I.C., Ponting, C.P. and Voet, T. Single-Cell Multiomics: Multiple Measurements from Single Cells. Trends Genet 2017;33(2):155–168.

McCarthy, D.J., et al. Scater: pre-processing, quality control, normalization and visualization of single-cell RNA-seq data in R. Bioinformatics 2017;33(8):1179–1186.

McGinnis, C.S., Murrow, L.M. and Gartner, Z.J. DoubletFinder: Doublet Detection in Single-Cell RNA Sequencing Data Using Artificial Nearest Neighbors. Cell Syst 2019;8(4):329–337 e324.

McGinnis, C.S., et al. MULTI-seq: sample multiplexing for single-cell RNA sequencing using lipid-tagged indices. Nat Methods 2019;16(7):619–626.

Pezzotti, N., et al. Approximated and User Steerable tSNE for Progressive Visual Analytics. IEEE Trans Vis Comput Graph 2017;23(7):1739–1752.

Picelli, S., et al. Full-length RNA-seq from single cells using Smart-seq2. Nat Protoc 2014;9(1):171–181.

Reinius, B. and Sandberg, R. Random monoallelic expression of autosomal genes: stochastic transcription and allele-level regulation. Nat Rev Genet 2015;16(11):653–664.

Risso, D., et al. A general and flexible method for signal extraction from single-cell RNA-seq data. Nat Commun 2018;9(1):284.

Sheng, K., et al. Effective detection of variation in single-cell transcriptomes using MATQ-seq. Nat Methods 2017;14(3):267–270.

Sheng, K. and Zong, C. Single-Cell RNA-Seq by Multiple Annealing and Tailing-Based Quantitative Single-Cell RNA-Seq (MATQ-Seq). Methods Mol Biol 2019;1979:57–71.

Stegle, O., Teichmann, S.A. and Marioni, J.C. Computational and analytical challenges in single-cell transcriptomics. Nat Rev Genet 2015;16(3):133–145.

Stoeckius, M., et al. Cell Hashing with barcoded antibodies enables multiplexing and doublet detection for single cell genomics. Genome Biol 2018;19(1):224.

Tang, F., et al. Deterministic and stochastic allele specific gene expression in single mouse blastomeres. PLoS One 2011;6(6):e21208.

Tukiainen, T., et al. Landscape of X chromosome inactivation across human tissues. Nature 2017;550(7675):244–248.

Usoskin, D., et al. Unbiased classification of sensory neuron types by large-scale single-cell RNA sequencing. Nat Neurosci 2015;18(1):145–153.

Van der Auwera, G.A., et al. From FastQ data to high confidence variant calls: the Genome Analysis Toolkit best practices pipeline. Curr Protoc Bioinformatics 2013;43:11 10 11–33.

Villani, A.C., et al. Single-cell RNA-seq reveals new types of human blood dendritic cells, monocytes, and progenitors. Science 2017;356(6335).

Wagner, J.M., et al. A comparative analysis of single cell and droplet-based FACS for improving production phenotypes: Riboflavin overproduction in Yarrowia lipolytica. Metab Eng 2018;47:346–356.

Wainer-Katsir, K. and Linial, M. Human genes escaping X-inactivation revealed by single cell expression data. BMC Genomics 2019;20(1):201.

Wolock, S.L., Lopez, R. and Klein, A.M. Scrublet: Computational Identification of Cell Doublets in Single-Cell Transcriptomic Data. Cell Syst 2019;8(4):281–291 e289.

Xin, Y., et al. Use of the Fluidigm C1 platform for RNA sequencing of single mouse pancreatic islet cells. Proc Natl Acad Sci U S A 2016;113(12):3293–3298.

Zeisel, A., et al. Brain structure. Cell types in the mouse cortex and hippocampus revealed by single-cell RNA-seq. Science 2015;347(6226):1138–1142.

Zhang, X., et al. Comparative Analysis of Droplet-Based Ultra-High-Throughput Single-Cell RNA-Seq Systems. Mol Cell 2019;73(1):130–142 e135.

Zheng, G.X., et al. Massively parallel digital transcriptional profiling of single cells. Nat Commun 2017;8:14049.

Zilionis, R., et al. Single-cell barcoding and sequencing using droplet microfluidics. Nat Protoc 2017;12(1):44–73.

